# SMC Motor Proteins Operate at the Near-Minimal Forces for DNA Loop Extrusion

**DOI:** 10.64898/2026.03.05.709531

**Authors:** Adrian John Pinto, Biswajit Pradhan, Damla Tetiker, Maurice Pierre Schmitt, Eugene Kim, Peter Virnau

## Abstract

Loop extrusion by structural maintenance of chromosomes (SMC) complexes is essential for genome organization, yet the forces driving this process remain poorly understood. We present a coarse-grained model enabling predictive simulations of *in vitro* loop extrusion experiments at experimentally relevant time and length scales by matching parameters with concrete experiments. Using this model, we demonstrate that the extrusion forces generated by SMC motor proteins are just sufficient to overcome initial entropic barriers and sustain loop extrusion, highlighting that motors operate in the thermal regime. By measuring stalling tension directly, we confirm that they can be reliably determined by the Marko-Siggia equation and that varying grafting distances in experimental setups has only a marginal effect on the resulting tension. These results provide a predictive computation method for dissecting the mechanics of SMC driven genome folding.

## 1 Introduction

The compaction of meter-long DNA into micron-scale cellular volumes represents a fundamental biological challenge that is mediated through multiple regulatory pathways. Among these, loop extrusion has emerged as a prominent mechanism for modulating DNA topology through the formation of chromatin loops. This three-dimensional organization of chromosomes into loops and topologically associating domains (TADs) constitutes a fundamental principle of genome architecture and function [1, 2, 3, 4, 5, 6, 7, 8, 9, 10]. Structural maintenance of chromosomes (SMC) complexes mediate this process, with their loop extrusion activity having been directly visualized through single-molecule *in vitro* imaging [11, 12, 13, 14, 15, 16, 17, 18, 19, 20].

Early theoretical frameworks demonstrated that progressive motors extruding along a polymer substrate could recapitulate key genomic features, including genome compaction and loop domain formation, typically through prescribed extrusion velocities or stochastic stepping rules [21, 22]. Subsequent three-dimensional polymer simulations expanded these approaches to investigate the roles of DNA topology, spatial confinement, supercoiling, and non-equilibrium driving forces in large-scale loop formation [23, 24, 25, 26, 27]. Recent advances have incorporated mechanical feedback mechanisms, revealing that DNA tension can modulate SMC translocation rates [28] and that directed extrusion may arise from anisotropic motor-polymer interactions [29]. Concurrently, experimental investigations have probed force-dependent extrusion dynamics under controlled tension and confinement conditions [12, 13], motivating the development of quantitatively calibrated models that integrate polymer topology, mechanics, and tension [30]. Despite these significant advances, a comprehensive framework that explicitly treats extrusion as a force-driven process while faithfully reproducing experimental geometries and stalling responses remains elusive.

Our modeling approach builds upon previous polymer-based loop extrusion simulations but introduces innovations in extrusion implementation, control mechanisms, and experimental calibration. While earlier work by Brackley et al. [25] demonstrated that passive slip-link sliding can in principle generate large loops, this mechanism is insufficient to account for loop formation in grafted chains as observed in experimental studies [12, 13]. Our model replaces the passive slip-link with an active motor, enabling loop growth through active extrusion while allowing tension-dependent stalling to emerge naturally from the underlying polymer mechanics. Although related force-sensitive models have demonstrated tension-dependent SMC translocation [28, 31], our work extends this perspective to match concrete experimental systems. This framework enables validation of traditional tension measurement approaches [32] and provides estimates of motor strengths from simulation data.

## 2 Results

We employ a modified version of a coarse-grained bead-spring model [33] capable of capturing the structural and topological properties of DNA [34, 35, 36]. To replicate experimental conditions, the two termini are grafted to a repulsive wall, and the chain is threaded through a handcuff structure composed of two rigid rings representing a SMC loop extrusion complex. In each time step of our molecular dynamics simulations, extrusion forces are applied to the monomers closest to the respective ring centers and perpendicular to the plane of the corresponding ring. Further details of the model are provided in Section 4. In the following section, we describe the calibration of forces and matching of time scales, which enable predictive simulations at experimentally relevant time and length scales.

### 2.1 Determination of force and time scale for quantitative simulations

The extrusion force serves as a tunable simulation parameter that governs both the extrusion rate and the maximum achievable loop size. For a given extrusion force, the non-extruded chain segments reach a characteristic steady-state extension that remains largely independent of the extrusion dynamics. To facilitate direct comparison with experimental observations, we define the relative extension that quantifies the stretching of the non-extruded linear segments:

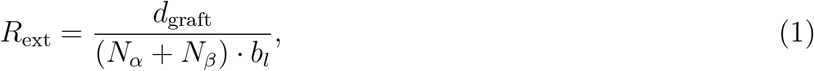

where *N*_*α*_ and *N*_*β*_ represent the number of monomers in the two non-extruded segments, respectively, *d*_graft_ denotes the grafting distance, and *b*_*l*_ the bond length. To establish correspondence between simulation parameters and experimental forces, we calibrate the extrusion force to achieve comparable maximum relative extensions to those observed experimentally. As demonstrated in section 2.2, the minimum force required for loop extrusion falls within the thermal regime. The maximum relative extension is determined for each independent loop extrusion event, as illustrated in Figure 1(a). (For conversion factors and further details, see section 4.) While experimental measurements exhibit substantial variability in maximum relative extension, simulations display a considerably narrower distribution for a given extrusion force. This discrepancy primarily stems from the absence of motor association/dissociation dynamics in our model. Since our investigation focuses on active extrusion dynamics rather than binding kinetics, we intentionally exclude the binding dynamics of the handcuff complex, which reduces variability-particularly by eliminating low-extension events. Furthermore, the direct tracking of individual beads in simulations yields significantly lower measurement uncertainty compared to experiments, where extensions are inferred from intensity measurements [13, 12, 11, 37].

**Figure 1.**
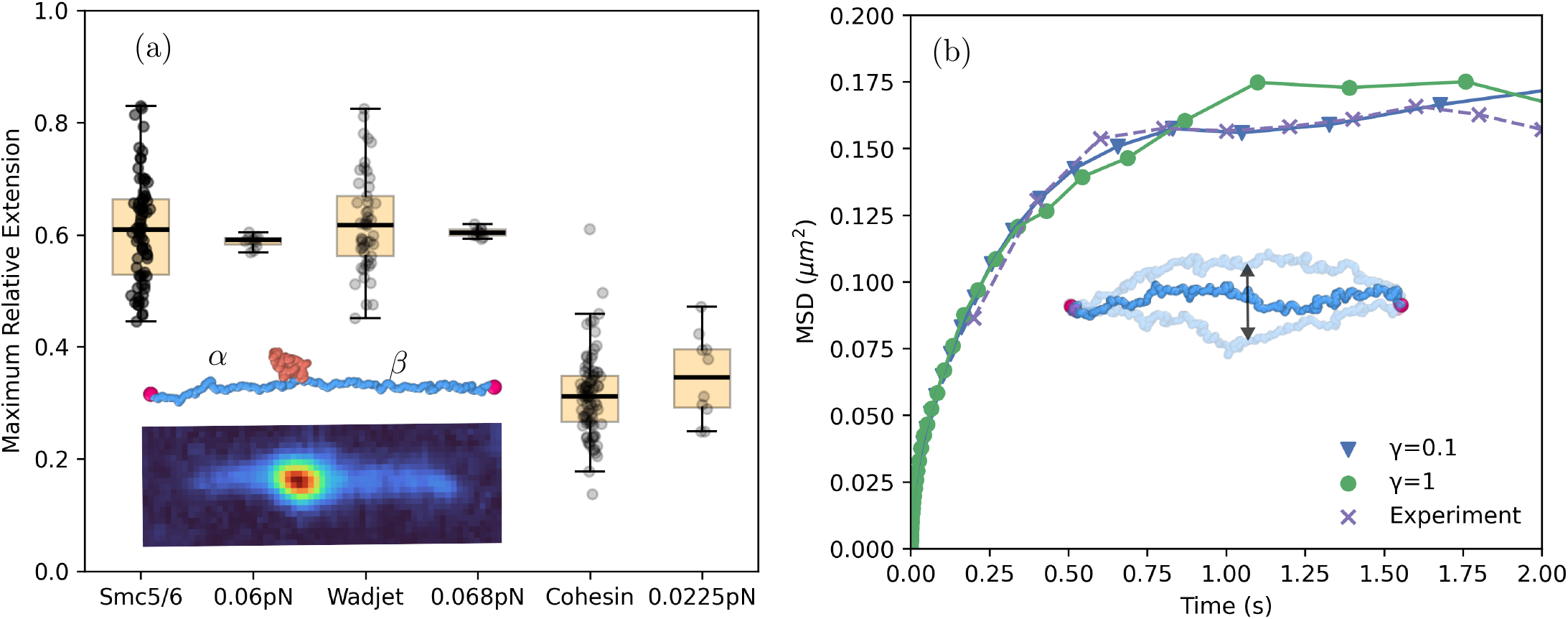
(a) Distributions of relative extension after loop extrusion for experiments (left) and simulations (right with force estimates) (*λ*-DNA, grafting distance: 5*μ*m for Wadjet, Smc5/6 and 3.8*μm* for cohesin) obtained by calculating the mean of the top 5% values of relative extension for each DNA looping event. The schematics show the completely extruded system in simulation and experiment; *α* and *β* denote the unextruded segments. Parts of this figure have been adapted from [37]. (b) Mean square displacement (MSD) of the segment located halfway between the grafted termini of the DNA in experiments and our simulation model for two different values of *γ*. The saturation of the MSD is matched to experiments to convert simulation time into real time.

Establishing correspondence between simulation and experimental time scales presents a more complex challenge. The system exhibits two relevant characteristic time scales: (a) a diffusive time scale, derived from the diffusion of the grafted polymer’s central segment, and (b) an extrusion time scale, determined by matching the initial extrusion rates between simulation and experiment. In an ideal scenario, both time scales would yield identical mapping parameters. However, our coarse-grained model implements continuous extrusion and does not account for extended waiting times observed experimentally between successive extrusion events. Incorporating such intermittent dynamics would significantly increase computational demands while offering limited additional mechanistic insight into the extrusion process. Therefore, we model extrusion as a continuous process to maintain computational efficiency while preserving the essential physical characteristics of the system.

To determine the diffusive time scale, we calculate the mean square displacement (MSD) of the polymer’s in-plane component perpendicular to the line connecting the grafted termini at half the grafting distance. Due to its geometric constraints, the MSD reaches a plateau at long times, as demonstrated in Figure 1(b). The Langevin thermostat’s friction coefficient *γ* serves as a temporal scaling parameter: increased *γ* slows the dynamics, while decreased *γ* accelerates them. When we normalize time scales between simulation and experiment by equating either the diffusion constants or, in our case, the time required to reach maximum lateral extension, simulations with different *γ* values collapse onto a common curve (Figure 1(b)) as explained in more detail in section 4. This scaling behavior validates our approach and enables substantial computational acceleration.

The extrusion time scale is established by matching the rate of change of the linear segments during the initial growth phase between experiments and simulation. We partition the polymer chain into three distinct regions: the extruded loop and the two unextruded linear segments, denoted as *α* and *β* (in Figure 1(a)). During this initial phase, the loop expands while the linear segments contract. Figure 2 presents the time derivatives of the linear segment sizes for both simulations (panel a) and experiments (panel b). The temporal conversion factor is systematically adjusted until optimal agreement between simulation and experimental data is achieved. Notably, when employing this extrusion time scale, the process occurs approximately two orders of magnitude faster than the diffusive time scale would suggest. As mentioned above, these two characteristic time scales could in principle be reconciled by incorporating waiting periods through temporary immobilization of monomers at the ring structures after an extrusion period.

**Figure 2.**
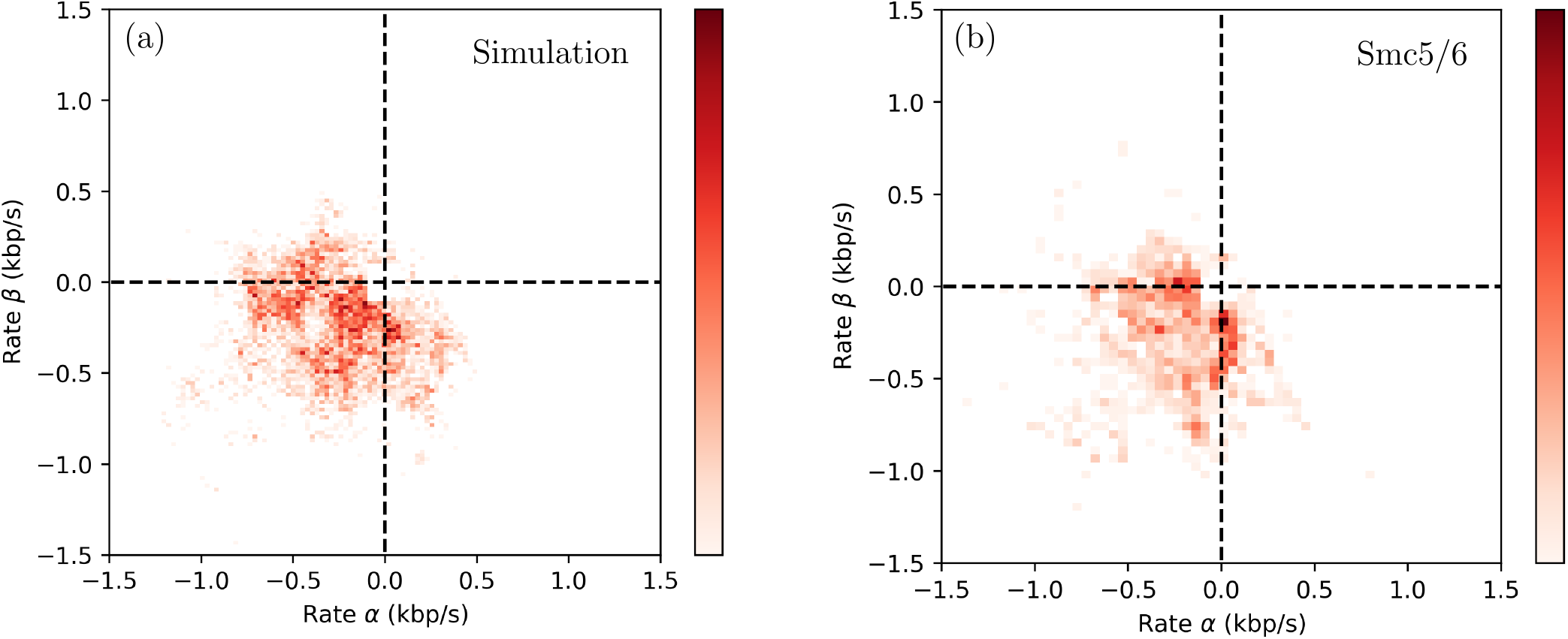
Rates at which the two linear DNA segments (*α* and *β*) DNA shrink during loop extrusion, in simulation for 0.06pN (a) and Smc5/6 experiments (b). The rates below |0.1| are omitted due to resolution limits in experiments. This figure has been adapted from [37].

### 2.2 Minimum motor forces for overcoming entropic barrier in loop extrusion simulations

One of our key findings is illustrated in Fig. 3, which highlights the efficiency of the loop extrusion process. The reduction in configurational entropy due to loop formation in polymers is a well-established concept in polymer physics, arising from the loss of accessible microstates imposed by topological constraints. When a grafted polymer is threaded through the two rings of the handcuff, a permanent topological constraint is introduced, ensuring the persistent existence of a loop. In this configuration, the minimum free energy state corresponds to the smallest possible loop size, as increasing the loop length further restricts the number of accessible polymer configurations. Consequently, enlarging the loop requires work to overcome an entropic free energy barrier, underscoring the necessity of an active extrusion force.

**Figure 3.**
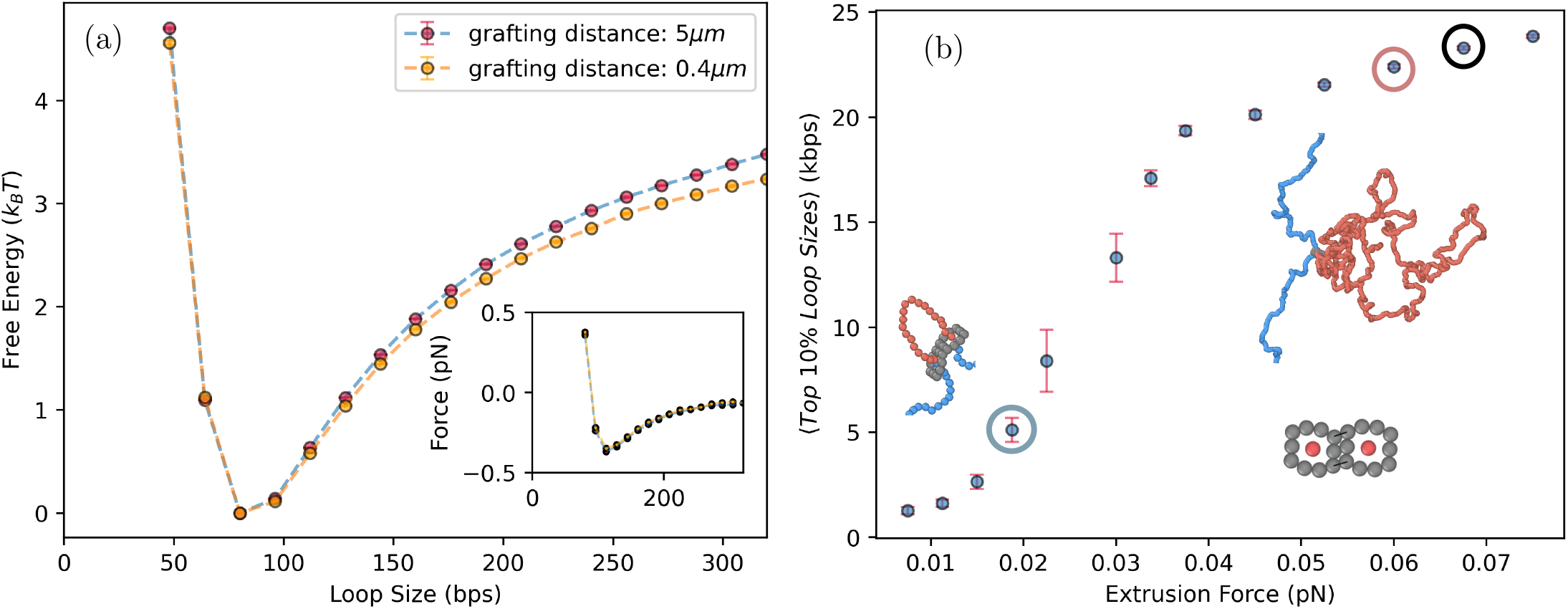
(a) Free energy (in units of *k*_*B*_*T*) as a function of the loop size derived from simulating a grafted *λ*-DNA chain threaded through a handcuff representing a SMC motor protein. The free energy was evaluated from the probability distribution of the loop size in the absence of extrusion forces. The free energy minimum does not occur at a loop size of zero due to steric hindrances from the handcuff, which enforce a finite loop size. The inset in figure (a) shows the derivative of the free energy with respect to the loop size. (b) Average of the largest 10% of loop sizes as a function of increasing extrusion forces (for a grafting distance of 5*μ*m). The loop size is averaged over a duration much greater than the duration needed for the loop to reach the mature phase. The schematics show the extrusion setup at low and high values of extrusion forces, and the handcuff structure. The red highlighted point is the force calibrated for the Smc5/6 complex, the black highlighted point depicts the force calibrated for the Wadjet complex and blue circle refers to cohesin. The forces correspond to the constant extrusion force in our simulation model which is required to obtain the same relative extension as in the experiments.

To quantify this barrier, we construct a free energy profile from the probability distribution of loop sizes obtained in simulations. Specifically, we thread the grafted polymer through the handcuff rings and perform simulations in the absence of any active extrusion forces. From these simulations, we compute the probability distribution *P* (*l*) of the loop size and define the corresponding free energy landscape as *F* (*l*) = *−k*_*B*_*T ln*(*P* (*l*)) (Fig. 3(a)). This profile exhibits a pronounced minimum at small loop sizes and depends only marginally on the grafting distance. The derivative of this free energy with respect to loop size, shown in the inset of Fig. 3(a), represents a restoring force that drives the system back toward the minimum loop size. This force is of the order of thermal fluctuations, indicating that thermal noise alone can induce transient small loops in the handcuff system but is insufficient to sustain larger loops. We determine the minimum extrusion force required to overcome this barrier and sustain extrusion in Fig. 3(b), which shows the average loop size as a function of the applied extrusion force. At low forces (*<* 0.02*pN*), loops remain transient and vanish upon time averaging. Once the extrusion force exceeds this threshold, stable loops emerge, leading to a sharp increase in the average loop size. For our system, which mimics Smc5/6, the extrusion force corresponds to 0.06 pN, indicating that this value is slightly above the threshold required to sustain loop extrusion. Similarly, for the Wadjet complex, the force is about 0.068pN. For comparison, we also considered cohesin, whose extrusion mechanism may or may not be symmetric [14, 7]. Interestingly, cohesin’s lower extrusion force, which is located right in the transition region, correlates with more frequent directional switching [37]. Although our model arguably simplifies the process by applying a constant force and modeling SMC complexes as a handcuff, this result nevertheless suggests that Smc5/6 and Wadjet operate within the thermal regime just above the required threshold, enabling them to fulfill their function without applying excessive forces or wasting energy.

### 2.3 Validation of the Marko-Siggia approach for determining stalling tensions

The Marko-Siggia equation is a widely used interpolation formula derived in 1995 [32] to describe the force-extension behavior of a semiflexible polymer, such as DNA, based on the worm-like chain (WLC) model. This equation effectively approximates the force required to extend a DNA chain of persistence length *l*_*p*_, which is grafted at one end and pulled at the other, to an extension *z/L* with *L* being the contour length (as illustrated in the upper snapshot of Figure 4(b)):

**Figure 4.**
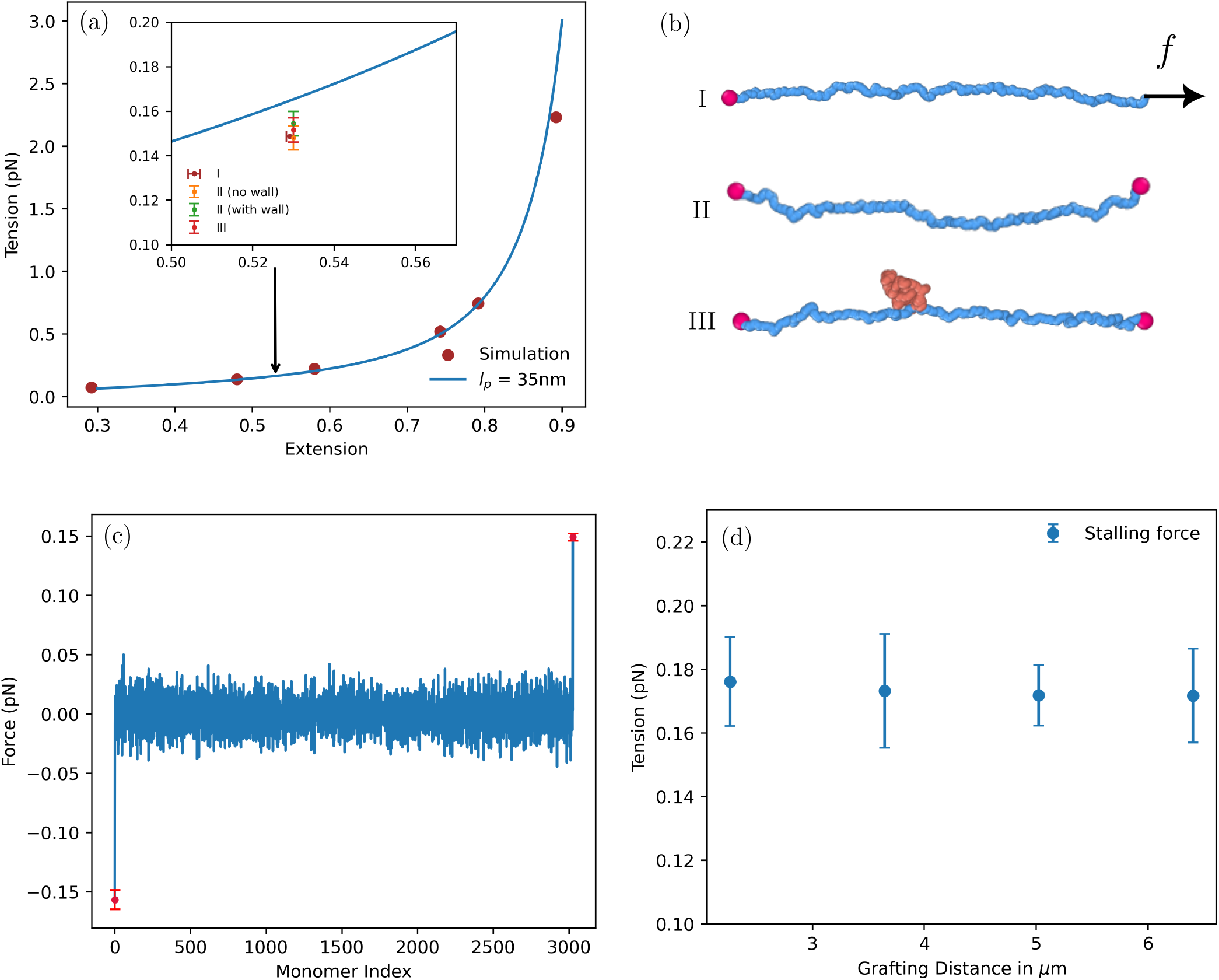
(a) Comparison of force-extension curves from our simulation model with the Marko-Siggia equation for persistence length *l*_*p*_=35*nm*. The inset compares simulation results for tension of a chain grafted on one side and pulled on the other I (Marko-Siggia scenario), a chain grafted at extension *z/L* with and without walls for which the loop has been removed II, a grafted chain with a fully extruded loop III. (b) Schematics of the simulation setup for force-extension measurement and tension measurements in the presence and absence of a loop. The beads are scaled up in size for visualization. (c) Tension profile measured directly from simulation by computing the z-component of force acting on the fixed beads at both the ends for the extruded system in the presence of walls. (d) Effect of grafting distance on the stalling tensions. The stalling tensions were evaluated using average relative extensions and equation (2) from 10 independent simulations.

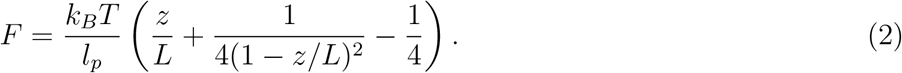

Figure 4(a) shows that forces applied to the free terminus of our polymer model agree well with equation (2). In the context of loop extrusion experiments, this formula is frequently employed to estimate stalling tensions. However, this application is based on several assumptions: First, it is presumed that the loop can effectively be removed from the full system (Figure 4(b) - III), and the stalling tension can be approximated by considering only the unextruded sections of the DNA. Then, it is assumed that grafting the chain at both ends to a specific extension *z* (Figure 4(b) - II) yields the same tension as pulling the chain with a corresponding force to achieve an average extension *z* (Figure 4(b) - I). Finally, interactions of the DNA with the wall, to which it is grafted to, should have no influence on tension.

To validate these assumptions, we conducted computer simulations tailored to our experimental setup. Specifically, we determined the average forces on the grafted termini along their connection line as exemplified in Figure 4(c) for the full system, which includes the extruded loop and wall interactions. The inset of Figure 4(a) provides a detailed comparison of the forces on the grafted beads for the reduced grafted chain with and without walls, alongside the full system, in relation to the Marko-Siggia equation and a chain pulled at one end. Our findings reveal that all scenarios yield tensions that are remarkably similar, differing by only a few percent from the values predicted by the Marko-Siggia equation. Notably, the extrusion force applied in each time step does not fully account for the measured tension in the system, implying the presence of additional entropic forces potentially arising from the loop. In Figure 4(d), we show that the stalling tension changes only marginally as long as the initial extension is significantly below the maximum possible extension for the motors.

## 3 Conclusion

In this study, we introduce a coarse-grained simulation model that facilitates direct quantitative comparisons with corresponding loop extrusion experiments. This progress is achieved through the model’s computational efficiency and parameter matching with concrete experiments, enabling simulations at experimental time and length scales. The full control over our simulations not only provides mechanistic insights into loop extrusion experiments but also allows for *in silico* modeling, offering a rapid and reliable means to explore potential application scenarios before conducting more complex experiments. Using this model, we demonstrate that extrusion forces from SMC motor proteins, which are on the order of thermal fluctuations, are just sufficient to overcome initial entropic barriers to loop extrusion and sustain the process. This barrier emerges because, from an entropic perspective, it is favorable to minimize the loop size to maximize configurational entropy for the remainder of the chain. Our simulations reveal that extrusion forces (≈0.05-0.1 pN) generated by Smc5/6 and Wadjet are large enough to surmount this initial barrier and maintain extrusion, underscoring the overall efficiency of the process. Even though these values were obtained using a coarse-grained model and by applying constant instead of stepwise forces, they nevertheless provide strong indication that SMC motors indeed operate in the thermal regime. For comparison, conventional motors such kinesin (5-7 pN), myosin (3-4 pN), dyenin (1-7 pN), and RNA polymerase (14-25 pN) stall at much higher forces. Such low forces for SMC complexes might explain their highly dynamic extrusion, slippage, and direction switching [38, 37]; enabling efficient search of regulatory factors such as CTCF. Notably, cohesin, which has a lower extrusion force than Wadjet and Smc5/6, also shows more frequent directional switching [37], consistent with a motor that operates closer to the entropic threshold and is therefore more susceptible to thermal perturbations. Similarly, RNA polymerase [39], telomeric Rap1 arrays [40] and R-loops [41] have all been shown to impede loop extrusion suggesting that a wide range of DNA-bound factors can serve as regulatory barriers. The low force budget seems not a limitation but a evolved feature that enables sensitive regulation of genome organization.

Furthermore, we show that the Marko-Siggia equation, commonly used to describe stalling tensions, aligns with the stalling tensions measured directly in our coarse-grained simulation within the experimentally relevant force regime, which justifies typical assumptions made in the process. Our model also demonstrates that stalling tensions are nearly independent of grafting distances in our specific experimental setup. Future work will utilize our model to investigate the details of the loop extrusion process under tension in environments that more closely mimic *in vivo* conditions, including the presence of nucleosomes and transcription factors.

## 4 Methods Section

### Simulation Model

The coarse-grained simulations of DNA are based on a standard bead-spring model [42, 43]. Beads are connected with FENE springs,

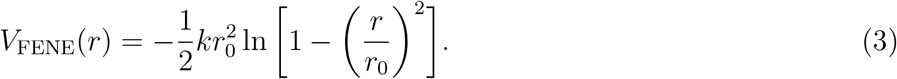

Chain stiffness is enforced with a Kratky-Porod potential with amplitude *κ*,

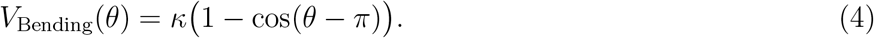

Non-bonded excluded volume interactions of the monomers are implemented via a Weeks-Chandler-Andersen potential,

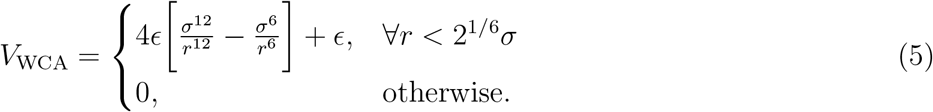

Just like in our experiments [12], the termini of the polymer are grafted onto a wall represented by a purely repulsive 9-3 Lennard-Jones potential,

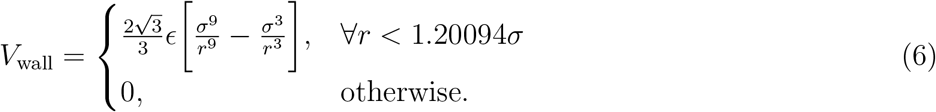

To map our experimental system onto the simulation model, we adapt our procedure from [33]. Screened charges on DNA are represented by effective excluded volume interactions following analytical approaches pioneered in the 1970s [44]. If charges are completely screened at high counter-ion concentrations, the size of a coarse-grained DNA bead corresponds to the locus of the DNA, i.e., 2.5 *nm*. In the partial screening scenario of our experiment (100 mM NaCl and 7.5 mM MgCl_2_) we set the bead size to 5.344 *nm* and equate this value with the expectation value of the bond potential at 0.9677*σ*, which enables investigations with molecular dynamics simulations. The persistence length of DNA is slightly reduced to 35 *nm* due to the presence of intercalating dyes [45], and the stiffness parameter *κ* ≈ 35*/*5.344 *k*_*B*_*T* following standard expressions for the worm-like chain model [46]. The number of beads *N* of the coarsegrained chain is given by equating the contour length of *λ*-DNA (16.167 *μm*) with the contour length of our model ((*N* − 1) * 5.344 *nm*) resulting in 3026 monomers. Likewise, the experimental grafting distance of 5*μm* corresponds to 909*σ*. The forces are measured in units of *k*_*B*_*T/σ* in simulations, which can be converted to experimental units (at 300K) by multiplying simulation units with a factor of 4.12*pNnm/*5.52*nm*. SMC motor proteins are represented by a simple handcuff consisting of two rings with 10 particles each (see depiction in Figure 3(b)) as suggested in [25]. Rings are modeled as rigid bodies and attached to each other with a FENE potential at two anchor points. A bending cost is introduced to each of the anchors to favor planar alignment. The total mass of the handcuff is set to be 5 times the mass of one monomer in the chain. While in [25] only the passive case was considered, here, we pull the DNA through the loop by applying extrusion forces to the two beads closest to the respective ring centers and equal and opposite forces to the rings. Even though we only consider two-sided extruders here, the model can in principle also be adapted to one-sided extrusion by applying asymmetric forces to the two central beads. Equations of motions were integrated using a standard Langevin thermostat with *dt* = 0.01*τ* and *γ* = 0.1 (see section 2). In our simulations, *γ* takes the role of a temporal scaling parameter. This can be quantitatively understood via the Einstein-Smoluchowski relation [47], which establishes that the product of a free particle’s diffusion constant *D* and friction coefficient *γ* equals *k*_*B*_*T*. Consequently, reducing *γ* by an order of magnitude increases the diffusion constant by the same factor, effectively accelerating the system dynamics. For *γ* = 0.1, 1s of real time corresponds to 0.9 × 10^6^ simulation times *τ* or 0.9 × 10^8^ time steps. All molecular dynamics simulations were performed with HOOMD-blue on CPUs and GPUs [48]. The visualization was done using ovito [49].

### Experimental Methods

The single-molecule experiments described here were performed as previously reported in detail [37]. Briefly, the wild-type cohesin, Wadjet, and octameric Smc5/6 complex was purified as described therein. Single-molecule visualization of DNA was carried out using a custom-built HiLo (highly inclined optical light sheet) microscope based on a Zeiss AxioVert200 body equipped with a 100×/1.46 NA oil-immersion TIRF objective. Excitation was done by 561 nm and 638 nm lasers, with fluorescence detected on a sCMOS camera (PCO Edge 4.2) after spectral filtering. All experiments were conducted at 30°C.

Flow cells were assembled using PEG-passivated glass slides functionalized with biotin-PEG to enable streptavidin-mediated surface immobilization of biotin-end-labelled *λ*-DNA (48.5 kb). SMC complexes (1 nM for Smc5/6, 100 pM for Wadjet, and 2nM for cohesin) was introduced in imaging buffer containing 40 mM Tris-HCl pH 7.5, NaCl (50 mM for cohesin, 100 mM for Smc5/6 and Wadjet), *MgCl*_2_ (7.5 mM for cohesin and Smc5/6, 10 mM for Wadjet), 0.5 mg/ml BSA, 1 mM TCEP, 2 mM ATP, 200 nM Sytox Orange, and an oxygen-scavenging system. Images were acquired at 100 ms exposure per frame for 1000 s.

Image analysis was performed using custom Python software. DNA loop sizes were quantified from kymographs by partitioning fluorescence intensity into loop (I_loop_), upstream (I_up_), and downstream (I_down_) regions, and multiplying intensity fractions by the known *λ*-DNA contour length (48.5 kb). Loop extrusion rates were estimated from linear fits to moving window of 5 s of the loop growth phase (initial 25 s). The relative DNA extension was calculated as the ratio of the measured end-to-end distance to the contour length of *λ*-DNA and converted to force via a force–extension calibration curve from magnetic tweezers measurements, enabling estimation of the stalling force at maximum loop size.

## Acknowledgements

This project was funded by SFB 1551 Project No. 464588647 of the DFG (Deutsche Forschungsgemeinschaft). The authors gratefully acknowledge the computing time granted on the supercomputer MOGON II and III at Johannes Gutenberg University Mainz as part of NHR South-West. BP acknowledge Hessian Ministry of Science and Art MSCA Grant for support. AP and PV would also like to thank Apratim Chatterji for fruitful discussions. LLMs have been used to improve the language and readability of the text.

## Author Contributions

AP and PV conceptualized the model. AP wrote the simulation code with inputs from MS. AP conducted the molecular dynamics simulation and wrote the first draft under supervision of PV. BP, EK conceptualized and conducted the single molecule experiments. BP, EK, DT analyzed the experimental data. All authors revised and edited the manuscript.

## Conflict of Interest

The authors declare that there is no conflict of interest.

## Table of Contents

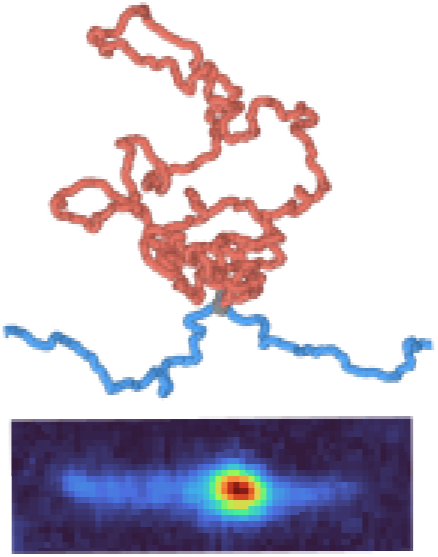

A coarse-grained model for DNA loop extrusion is developed using a force-based approach that quantitatively reproduces experimental stalling behavior, rate decays, and observed loop geometries. The framework enables simulation at experimentally relevant time and length scales, providing mechanistic insight into SMC motors. The model further allows validation of commonly used tension measurement approaches.

## Notes

### Competing Interest Statement

The authors have declared no competing interest.

### Summary of Updates

Removed trademark logos from the manuscript.

